# Molecular basis of inhibition of the amino acid transporter B^0^AT1 (SLC6A19)

**DOI:** 10.1101/2024.02.18.580905

**Authors:** Junyang Xu, Ziwei Hu, Lu Dai, Aditya Yadav, Yashan Jiang, Angelika Bröer, Michael Gardiner, Malcolm McLeod, Renhong Yan, Stefan Bröer

**Author notes:** Materials & Correspondence: Stefan Bröer or Renhong Yan. The authors contributed equally to the work (Junyang Xu, Ziwei Hu, Lu Dai, Aditya Yadav, Yashan Jiang).

## Abstract

The epithelial neutral amino acid transporter B^0^AT1 (SLC6A19) is the major transporter for the absorption of neutral amino acids in the intestine and their reabsorption in the kidney. Mouse models have demonstrated that lack of B^0^AT1 can normalize elevated plasma amino acids in rare disorders of amino acid metabolism such as phenylketonuria and urea-cycle disorders. This discovery has resulted in a quest to identify high-affinity selective inhibitors of B^0^AT1 for the treatment of disorders of amino acid metabolism. Here we employed a medicinal chemistry approach to improve lead compounds that were identified in a previous high-throughput screen. This effort yielded a group of compounds that inhibited B^0^AT1 with IC_50_-values of 31-90 nM. High-resolution cryo-EM structures of B^0^AT1 in the presence of two compounds from this series identified an allosteric binding site in the vestibule of the transporter. Mechanistically, binding of these inhibitors prevents a substantial movement of TM1 and TM6 that is required for the transporter to make a conformational change from an outward open state to the occluded state.

---

Amino acid homeostasis is an important factor in cellular and organismal physiology ^1^. Plasma amino acid levels are maintained in a narrow range and are used as a diagnostic tool to identify rare diseases, such as phenylketonuria, urea cycle disorders, tyrosinemia, and lysinuric protein intolerance ^2^. Smaller deviations of amino acid homeostasis occur in type 2 diabetes and are related to insulin resistance ^3,4^. Various strategies are used to treat derangements of amino acid metabolism, such as diet, nitrogen scavengers and supplements ^2^. A more recent strategy is to counteract elevated levels of amino acids by decreasing the uptake or increasing the loss of amino acids via inhibition of amino acid transporters.

Epithelial amino acid transporters are critical for the absorption of amino acids in the intestine and the reabsorption of amino acids in the kidney ^1^. Blockade or lack of these transporters causes reduced absorption in the intestine and loss of amino acids in the kidney ^5^. In humans, mutations in the *SLC6A19* gene cause Hartnup disorder, a largely benign condition characterized by aminoaciduria ^6^. Consistently, lack of the apical neutral amino acid transporter B^0^AT1 (Broad neutral amino acid transporter 1, encoded by the *SLC6A19* gene) has been shown to counteract elevated amino acids in phenylketonuria and urea cycle disorder mouse models ^7,8^. Moreover, lack of B^0^AT1 has also been shown to improve glucose tolerance through a variety of mechanisms including elevated levels of FGF21, GLP-1 ^9^ and reducing liver triglycerides ^10^ and may protect against kidney injury ^11^. As a result, significant efforts are underway to develop high-affinity selective inhibitors of B^0^AT1 ^10,12-16^. This has been accompanied by the development of variety of assays to measure B^0^AT1 activity, such as proteoliposomes ^12^, solid supported membrane electrophysiology ^16^, voltage-sensitive fluorescent dyes (FLIPR) ^14,15^ and classical radioactive flux assays ^14,15^.

The amino acid transporter B^0^AT1 belongs to the amino acid transporter subfamily of the SLC6 family ^17^. This subfamily is characterized by an extended loop linking transmembrane helices 7 and 8. In the case of B^0^AT1 this loop is required to form contacts with the single transmembrane helix (TM) proteins collectrin or ACE2, enabling cell surface expression in the kidney ^18^ and intestine ^19^, respectively. Mammalian cell lines overexpressing B^0^AT1 and collectrin have been employed to characterize inhibitors, but the endogenous transport activity in radioactive uptake assays is substantial, reducing the signal-to-noise ratio ^14^.

The structure of the B^0^AT1/ACE2 complex was recently resolved by single-particle cryo-EM ^20^. Despite the significant pharmaceutical interest in B^0^AT1 inhibitors, a structural understanding of their binding to the transporter is missing. In contrast, the pharmacology of neurotransmitter transporters of the SLC6 family is well developed. Most neurotransmitter transporter inhibitors act by binding to the orthosteric binding site (S1) but some also bind to an allosteric site in the vestibule of the transporters (S2) ^21-24^. Importantly, all serotonin reuptake inhibitors bind with higher affinity to the S1 site than to the S2 site ^24^.

Initially identified inhibitors of B^0^AT1 showed IC_50_ values in the 10-100 µM range ^12,14^, whereas inhibitors derived from high-throughput screening showed IC_50_-values in the range from 0.4-15 µM ^10,15^. A medicinal chemistry approach subsequently improved the IC_50_ of nimesulide analogues to compound 39 with an IC_50_ of 0.035 µM ^13^.

In this study we have improved the radioactive flux assay by suppressing endogenous transporters and used this improved assay to identify homologues of the recently identified B^0^AT1 inhibitor E4 (2-(4-chloro-2,6-dimethylphenoxy)-*N*-propan-2-ylacetamide). This resulted in improved compounds with nanomolar affinity. Structural analysis showed binding of these compounds to an allosteric site of the transporter. These results pave the way for rational drug design for an important pharmaceutical target.

## Results

To characterize novel inhibitors of B^0^AT1 we have used two independent assays, namely a commercial fluorescent plate-reader based assay (FLIPR) and a conventional radioactive uptake assay ^14^. While the FLIPR assay is suitable for high-throughput screening, detailed characterization of initial hits requires a secondary assay that directly measures amino acid uptake. Both assays are used in conjunction with a Chinese hamster ovary cell line stably overexpressing B^0^AT1 and collectrin (CHO-BC) ^14^ with leucine as a preferred substrate. However, CHO cells also have endogenous transporters for leucine ^25^. To identify the fraction of leucine uptake mediated by B^0^AT1, uptake is measured in the presence and absence of Na^+^ because B^0^AT1 is Na^+^-dependent, while the main endogenous leucine transporter LAT1 (SLC1A5) is Na^+^-independent ^14^ (Fig. 1A). Two problems arise from endogenous amino acid transport when characterizing B^0^AT1 inhibitors. First, inhibition can remain incomplete, even when Na^+^-independent transport via LAT1 is subtracted, due to the activity of other transporters. Secondly, inhibition of B^0^AT1 can result in an apparent increase of Na^+^-independent transport activity (LAT1), instead of an expected inhibition of Na^+^-dependent leucine transport (B^0^AT1) (Fig. 1A). The net uptake activity is reduced but for an optimal assay it would be better to isolate the B^0^AT1 component. Leucine uptake in CHO-BC cells is comprised of five components, namely B^0^AT1, LAT1 (SLC7A5), ASCT2 (SLC1A5) and SNAT2 (SLC38A2) ^26,27^ (Fig. 1B). However, SNAT2 activity can be suppressed by a recent change of media ^28^.

**Fig. 1:**
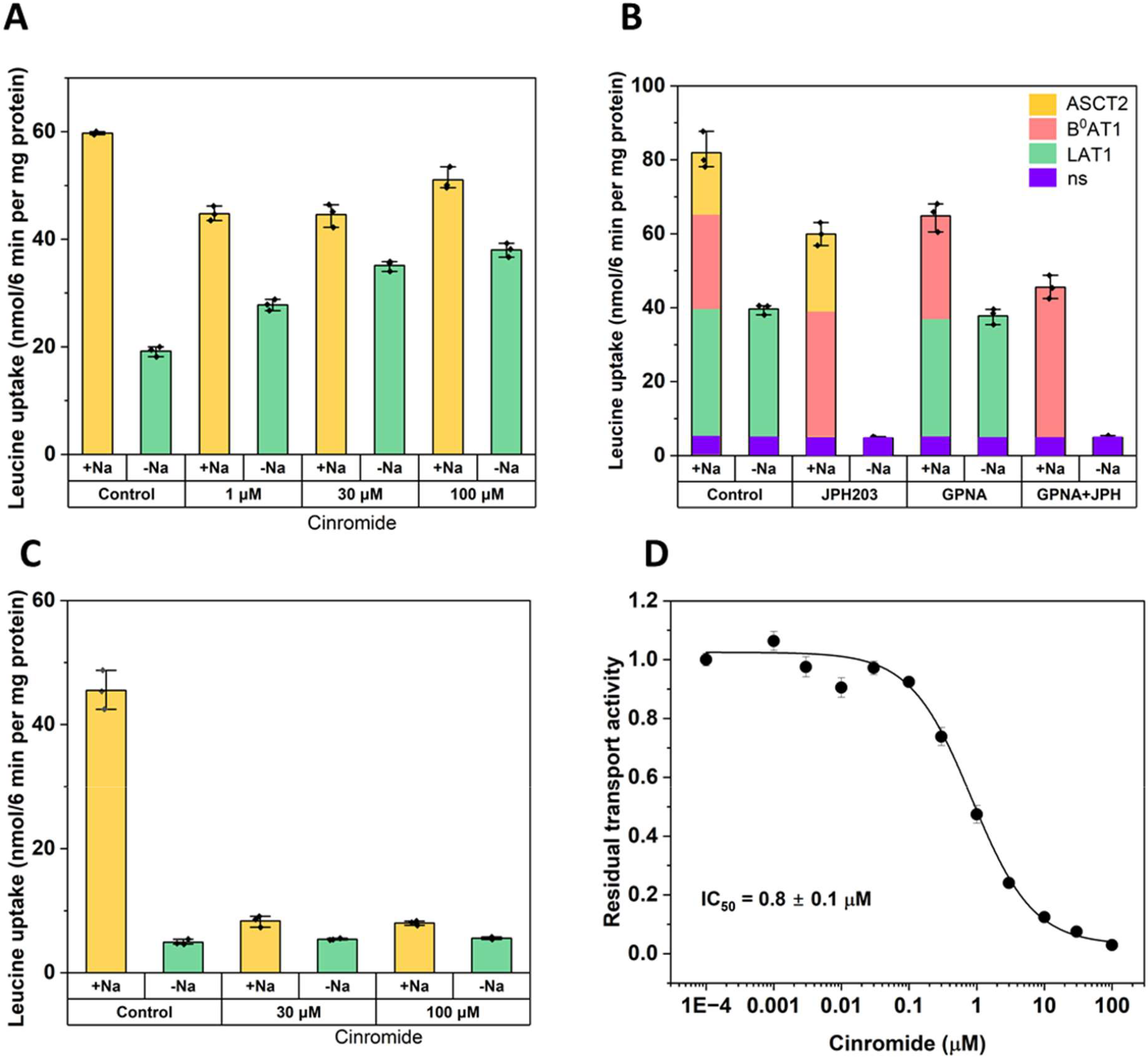
Endogenous contributions to leucine transport in the B^0^AT1 flux assay. Leucine transport was characterized in CHP-BC cells using uptake of [^14^C]leucine (n=3). A) Na^+^-dependent and Na^+^-independent components of Leucine uptake in CHO-BC cells. An apparent increase of Na^+^-independent leucine transport is observed after application of increasing concentrations of B^0^AT1 inhibitor cinromide. Leucine transport is blocked incompletely due to endogenous transport activities. B) The Na^+^-independent endogenous transport can be blocked by LAT1 inhibitor JPH203. The Na^+^-dependent component of leucine uptake that is sensitive to inhibition by GPNA was assigned to ASCT2. The remaining transport activity was assigned to B^0^AT1. Transport components are indicated by colour. C) In the presence of GPNA and JPH203, the remaining Na^+^-dependent transport activity is completely blocked by cinromide D) IC50 of cinromide as determined with the optimized assay. All data shown as mean ± SD, individual datapoints were overlayed in all bar graphs.

As a first step to improve the assay, we added JPH203 (3 µM), a high-affinity LAT1 inhibitor that blocks Na^+^-independent leucine transport without affecting B^0^AT1 ^29^ (Fig. 1B). The activity of ASCT2 could be suppressed by γ-glutamyl-p-nitroanilide (GPNA, 3 mM) ^30^. When both inhibitors were combined, the known B^0^AT1 inhibitor cinromide completely blocked the remaining transport activity without increasing Na^+^-independent transport (Fig. 1C). Using this optimized assay, we determined an IC_50_ value of 0.8 ± 0.1 μM for cinromide (Fig. 1D), which agrees with earlier measurements ^10,15^. An additional advantage of the optimized assay is the reduction of samples, because measurements can be performed in the presence of Na^+^ without the need to subtract endogenous transport activity in Na^+^-free buffer.

In a previous high-throughput screen we identified compound E4 (2-(4-chloro-2,6-dimethylphenoxy)-*N*-isopropylacetamide) as a high-affinity inhibitor of B^0^AT1 ^10^ with an IC_50_ = 7.7 ± 1.9 µM. When we ordered a new batch of the compound, we noticed that the inhibitory activity was strongly reduced such that an IC_50_ could not be determined (16% inhibition at 30 µM). Subsequent structural characterisation of the original and new batch revealed that the active compound was in fact (2-(4-chloro-3,5-dimethylphenoxy)-*N*-isopropylacetamide). We called this compound JX98 (Fig. 2) and refer to E4 as JX8 to avoid confusion and use these numbers from hereon. This revealed a strong structure-activity relationship (SAR), namely that the methyl-groups of the phenoxy moiety cannot be in the ortho-position but must be in meta-position (Fig. 2 and Table S1 (JX8 vs. JX98)). Even a single methyl-group in this position reduces the inhibitory potential to a similar extent (JX42). As a result, we performed all subsequent SAR experiments with JX98.

**Fig. 2.**
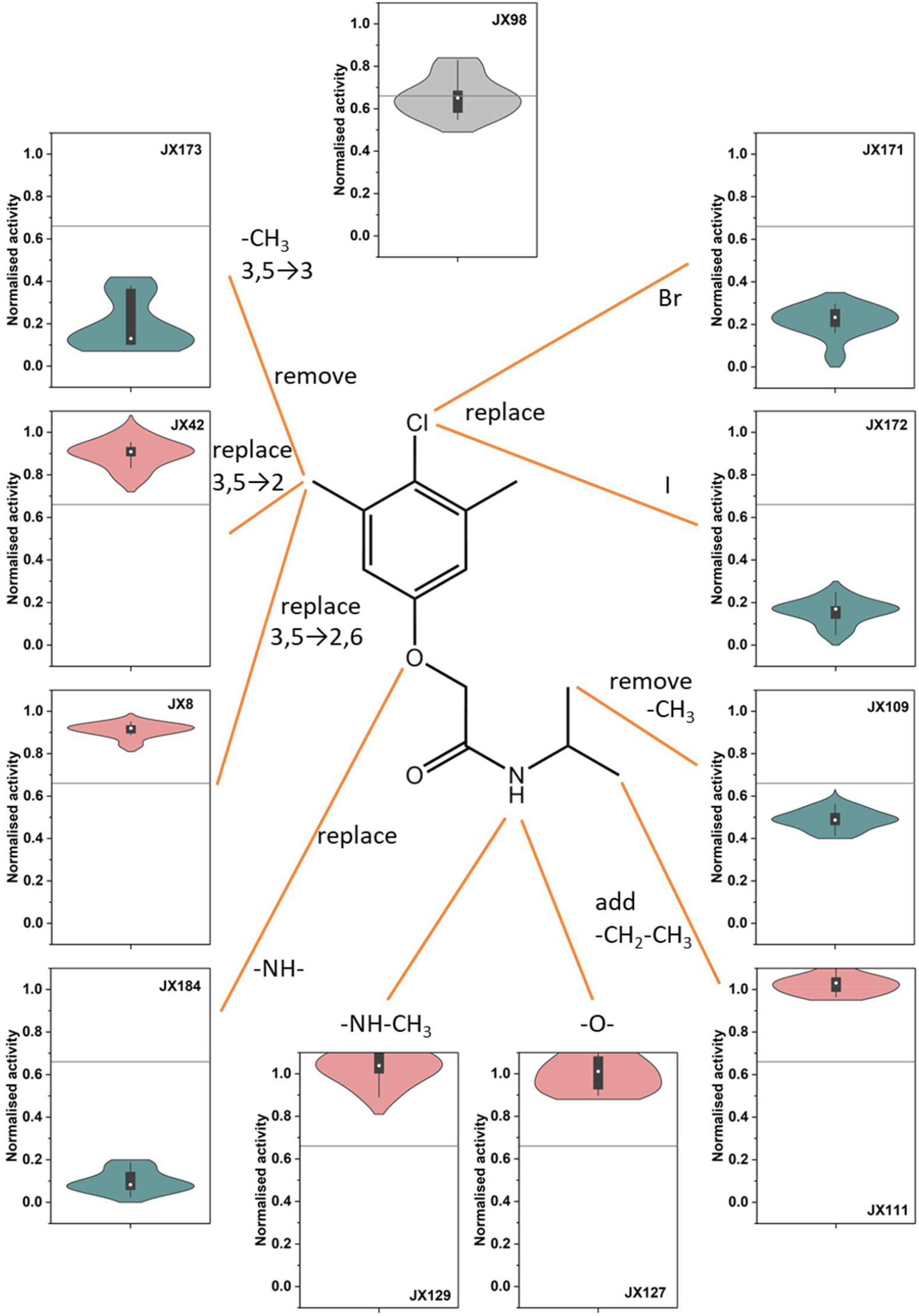
Structure-activity relationships of SLC6A19 inhibitor JX98. For each chemical modification the remaining transport activity in the presence of 30 μM inhibitor is shown as a violin plot (n=10) with median, SD, minimum and maximum indicated. Better inhibitors result in lower residual activity. Green violins represent improvements, red violins undesirable modifications. More details can be found in supplementary data.

The systematic variation of the JX98 pharmacophor is summarized in Fig. 2 with more detailed information presented in Table S1. Notably, the secondary amide of the sidechain was essential for inhibitory action and could not be replaced by an ester (JX127), thioester (JX128), *N*-methylated tertiary amide (JX129) or ketone (JX130) (Fig. 2 and Table S1). This suggests that a hydrogen bond donor may be required at this position. The X-ray crystal structure of JX98 (CCDC-2300201, Fig. S1) revealed intramolecular hydrogen bonding between the amide nitrogen and the phenolic ether ^31^, while the structure of JX225 (CCDC-2300200, Fig. S1) revealed intermolecular hydrogen bonding between the amide N-H and carbonyl of adjacent molecules, highlighting the potential of this secondary amide to mediate intramolecular or intermolecular contacts.

The improving features were combined and reiterated in a series of new compounds (JX225-228 and JX235-240). These yielded potent inhibitors with IC_50_-values ranging from 31 nM to 90 nM as measured using the FLIPR assay (Fig. 3). The corresponding IC_50_-values measured by radioactive uptake were higher, ranging from 111 nM to 440 nM.

**Fig. 3.**
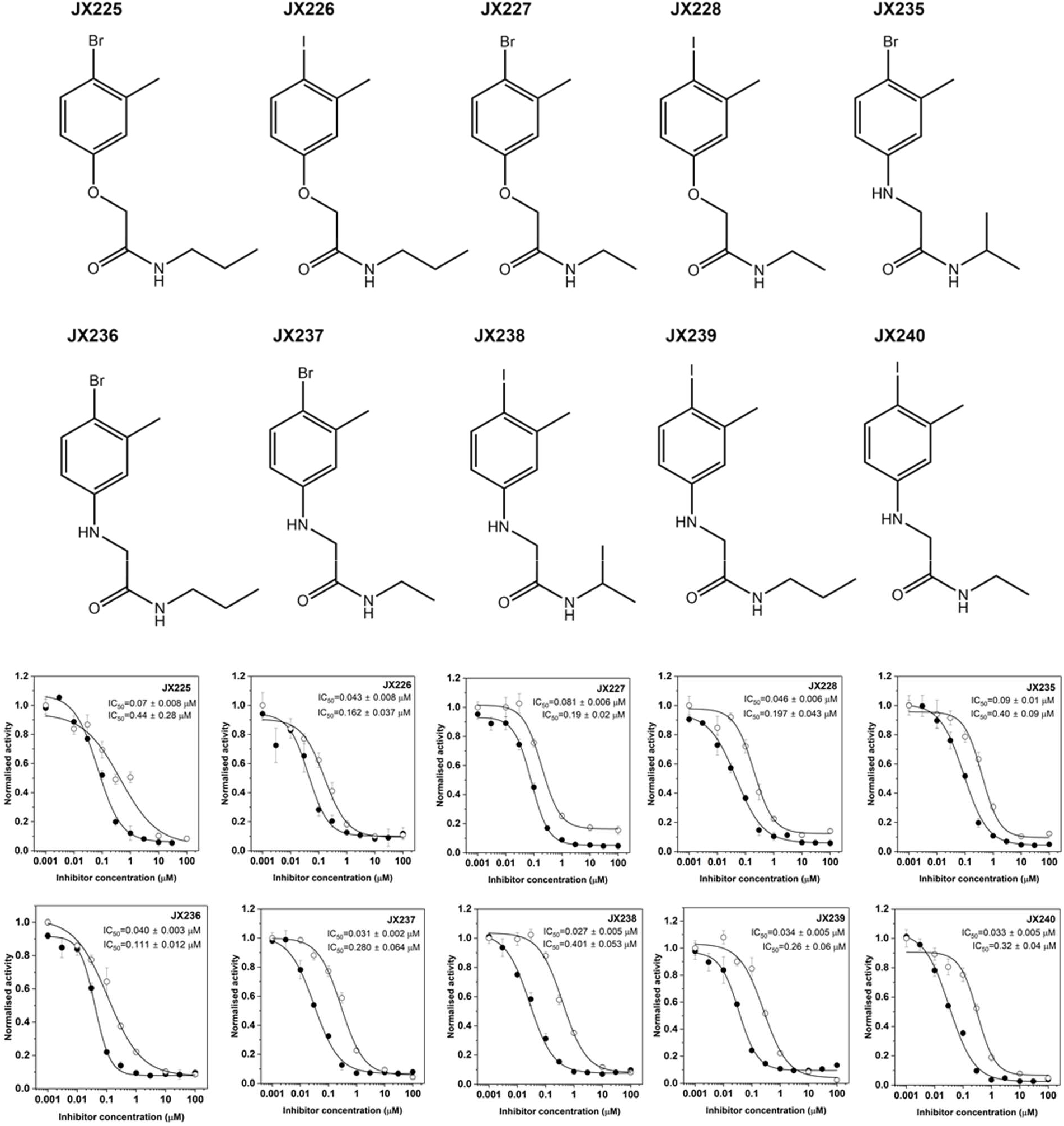
Pharmacological properties of optimized B^0^AT1 inhibitors. Structures of the optimized inhibitors are shown above. Leucine transport activity was determined by FLIPR assay (closed symbols) and by radioactive leucine uptake assay (open symbols) in CHO-BC cells at the indicated inhibitor concentrations (n=6). Data are shown as mean ± SD.

To understand the mechanism of inhibition further we determined the structure of B^0^AT1 in the presence of JX98 and JX225. The structures of ACE2-B^0^AT1 incubated with JX98 and JX225 were resolved at a remarkable resolution of up to 3.12 Å or 3.30 Å, respectively (Fig. 4A and Fig. S2-3 and Table S2). Detailed information regarding cryo-EM sample preparation, data collection, processing, and model construction can be found in the “Materials and Methods” section. Electron density maps for selected transmembrane helices are shown in Fig. S4.

**Figure 4.**
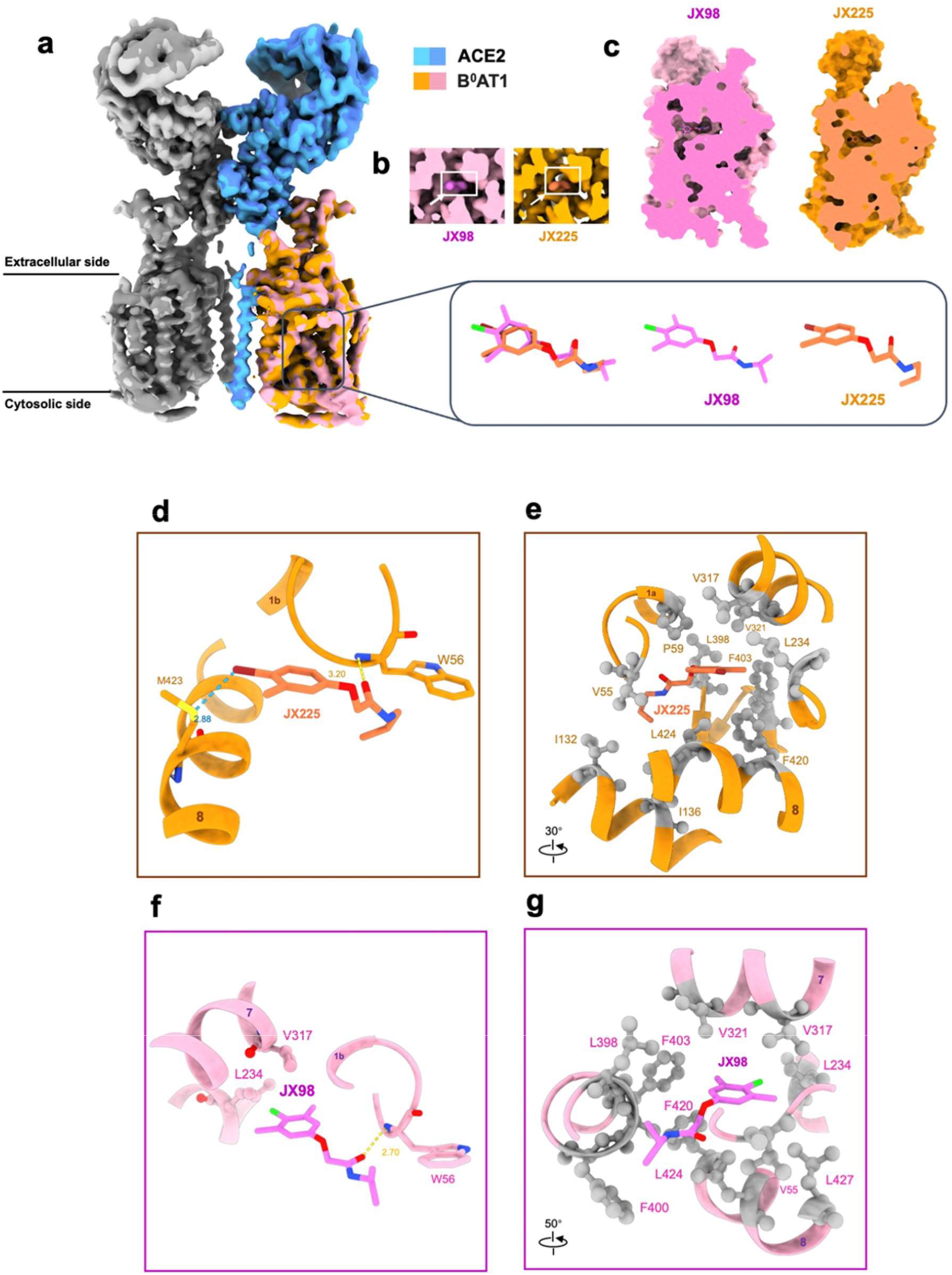
Overall structure of the ACE2-B^0^AT1 bound with inhibitors. (a) Cryo-EM map of the full length of the ACE2-B^0^AT1 heterodimer in complex with inhibitors. ACE2 and B^0^AT1 bound with JX225 (coral) are represented in sky blue and orange, respectively, while ACE2 and B^0^AT1 bound with JX98 (magenta) are shown in darker blue and pink, respectively. The structural comparison of JX-225 and JX98 is presented in the block. **(b)** The inserts in Cryo-EM map of the human ACE2-B^0^AT1 complex. The density corresponding to JX98 and JX-225 is color-coded in magenta and coral, respectively. **(c)** Positioning of JX98 and JX-225 in ACE2-B^0^AT1 complex. Both JX98 and JX-225 are centrally located within the presumed transport pathway. **(d, e)** The binding mode of JX-225 in B^0^AT1 is shown, highlighting hydrogen, halogen and hydrophobic interactions. **(f, g)** The binding mode of JX98 in B^0^AT1 is presented, with schematic diagrams illustrating the interaction environment, following the same representation as in (d, e).

Compared with the apo state of the ACE2-B^0^AT1 complex (PDB: 6M18), an extra density was found enclosed by TM1b, TM7, TM8 and EL_7-8_ in the vestibule of the transporter for both structures (Fig. 4B-C and Fig. S4). The high-resolution structures enabled model building of JX98 and JX225 molecules, which are both enclosed by a hydrophobic pocket (Fig. 4D-G and Fig. S5). As shown in figure 4, the structures of JX225 and JX98 exhibit remarkable similarity, as both occupy nearly identical positions within the same binding locale of B^0^AT1. Consistent with the analysis of structure-activity relationships, the backbone nitrogen of Trp56 on TM1 has the potential to form hydrogen bonds with the carbonyl oxygen atoms of JX225 and JX98, respectively (Fig. 4D and F), but there was no hydrogen bond to the secondary amine. The halogen atoms of both, JX225 and JX98 exert substantial binding forces, albeit through distinct mechanisms. Through model building, it was discerned that the role of halogen atoms is not uniform, indicating distinct modes of action for these atoms. The bromine atom of JX225 has the capability to establish a non-covalent halogen bond ^32^ with Met423 on TM8 although the angle is not optimal. This interaction may contribute to the increased affinity between JX225 and B^0^AT1 ^33^. In contrast, the chlorine atom of JX98 is located at greater distance from Met423 and engages in hydrophobic interactions with Val317 and Leu234 on TM7. This difference might be caused by the steric hindrance of methyl groups adjacent to the chlorine atom of JX98. The aromatic rings of both JX225 and JX98 exert strong hydrophobic interaction with B^0^AT1 including a series of residues (Fig. 4D-G). The hydrophobic environment surrounding JX225 comprises amino acids Val55, Pro59, Leu132, Leu136, Leu234, Val317, Val321, Phe403, Phe420 and Leu424. The hydrophobic environment surrounding JX98 is primarily mediated by residues Val55, Pro59, Val317, Leu398, Phe400, Phe403, Phe420, Leu 424 and Leu427 (Fig. S5).

Importantly, the JX98 and JX225 binding site is not located at the traditional substrate binding site of LeuT-fold transporters but rather overlaps with the S2 allosteric binding site identified in several transporters in this family ^24^. However, the allosteric binding of both inhibitors prevents a substantial movement in TM1 and TM6 of B^0^AT1 that is required for the transporter to make a conformational change from an outward open state to the occluded state (Fig. 5a,b). Notably, the side-chain of Trp56 has to rotate out of the way to allow binding of the inhibitors. To understand the role of Trp56, we mutated it to glycine and leucine (Fig. 5c). Mutation to glycine completely abolished transport activity, even below the level of ACE2 alone, which is most likely caused by preferential association of ACE2 with the heterologously expressed B^0^AT1. Mutation to leucine recovered some transport activity. While the transport activity of Trp56Gly was too low to assess inhibition by JX225, the residual transport activity of Trp56Leu appeared to be resistant to inhibition by JX225 (Fig. 5d). These results suggest that flexibility and size of the Trp56 side-chain plays a key role in the turnover of the transporter.

**Figure 5.**
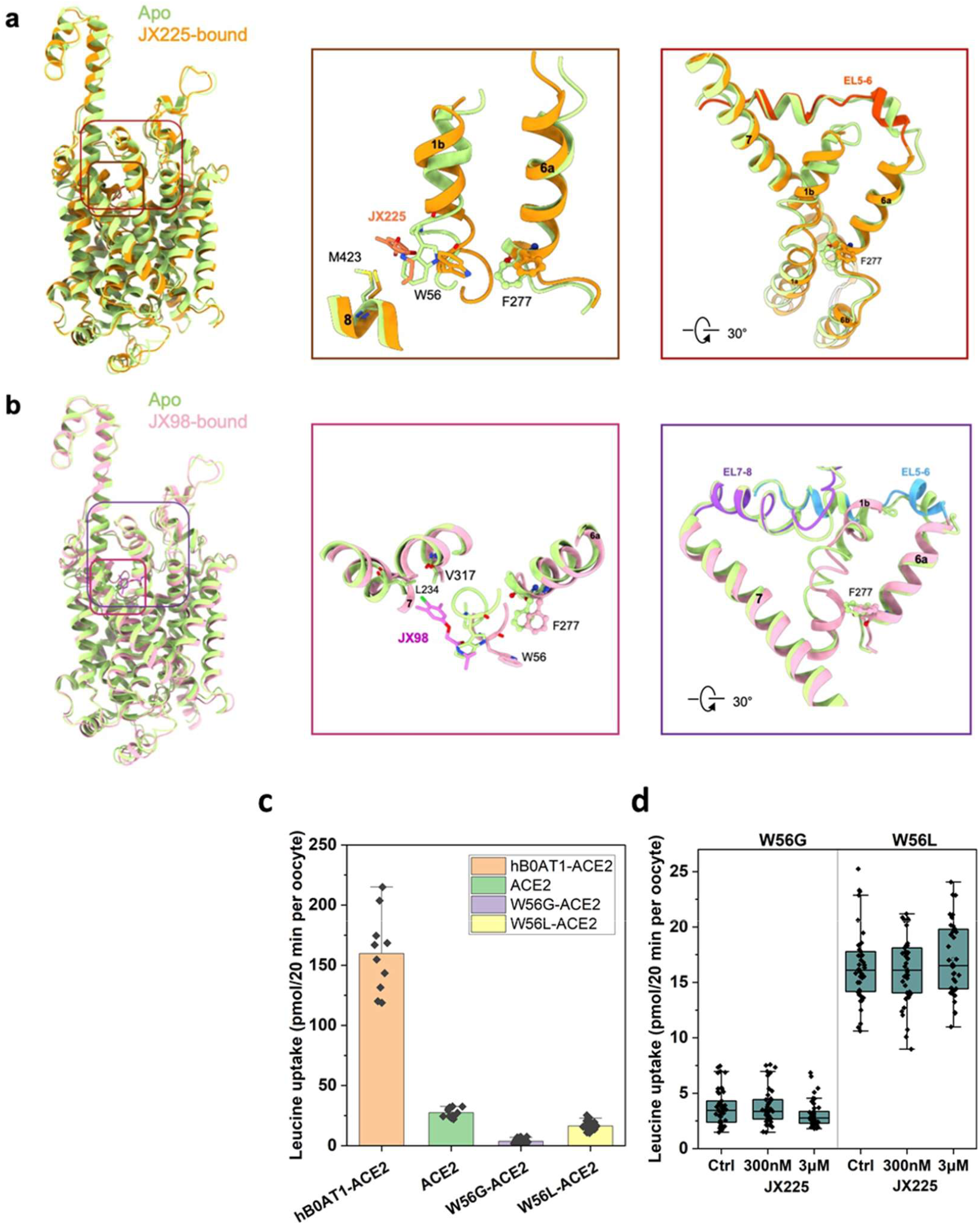
The movement of TM1 and TM6 and the role Trp56. **(a-b)** A comparison is drawn between the apo structure of B^0^AT1 (PDB: 6M18) when bound with JX225 and JX98, respectively. The apo structure, B^0^AT1 bound with JX225, B^0^AT1 bound with JX98 are coloured by green, orange and pink, respectively. The alterations in the positions of TM1 and TM6 are depicted in the middle and right panels. JX225 **(a)** and JX98 **(b)** are represented in coral and magenta, respectively. **(c)** *Xenopus laevis* oocytes were injected with 10 ng cRNA encoding B^0^AT1 (or its mutants W56G and W56L) and ACE2. Leucine uptake was measured after 4 days of expression **(d)** Leucine uptake was measured as in panel c in the absence and presence of inhibitor JX225 at the indicated concentrations. Bar graphs show mean ± SD and individual data points.

## Discussion

This study makes three distinct advances in the generation of selective and potent inhibitors of the pharmaceutical target B^0^AT1. First, an improved assay for characterization of inhibitors was developed, secondly, a high-affinity allosteric inhibitor of B^0^AT1 was generated, and thirdly, an allosteric inhibitory binding site was identified in the vestibule of B^0^AT1.

The first clinical trials with B^0^AT1 inhibitors for the treatment of phenylketonuria have been initiated (NCT05781399) after posting promising preclinical data ^34^. This demonstrates B^0^AT1 as a validated target for the treatment of orphan diseases with an overabundance of neutral amino acids. This could include orphan diseases such as alkaptonuria, urea cycle disorders and disorders of branched-chain amino acid metabolism. As a result, a better understanding of potential drug binding sites in members of the SLC6 amino acid transporter subfamily is important. In contrast to drugs that block serotonin, dopamine and noradrenaline transporters, which typically bind to the orthostatic S1 site ^23^, JX98 and JX225 bind to the allosteric S2 site. This is likely the result of the screening strategy employed to isolate the original lead compound E4 (JX98), which made use of an elevated concentration (1.5 mM) of substrate isoleucine against 10 μM inhibitor ^10^. This generates a significant competitive pressure excluding compounds that preferentially bind to the S1 site. Citalopram, which has been shown to bind to the S2 site in serotonin transporter SERT, still binds with 1000-fold higher affinity to the S1 site ^35^. Only recently has a preferential allosteric inhibitor for SERT being identified from a selection of citalopram analogues ^36^ Assignment to the S2 site was based on displacement experiments with labelled imipramine and citalopram. Here we provide direct structural evidence for selective binding to an allosteric site in B^0^AT1. While initial docking studies suggest that JX98 could bind to the S1 site ^10^, we were not able to find structures with an inhibitor bound to this site. The proposed binding pose at the allosteric site does not provide a hydrogen bond to the secondary amine, a feature we would have expected from the SAR. As a result, we cannot exclude that the inhibitor may also bind to the S1 site, where a hydrogen-bond could be formed. Notably, JX225 is structurally similar to the established B^0^AT1 inhibitor cinromide ^15^. Our results therefore suggest a similar binding mode for this compound.

The structure of the LeuT-fold amino acid:sodium symporter MhsT has been determined in an inward-facing conformation ^37^. In this structure the vestibule is collapsed, with Trp33, which is homologous to Trp56 in B^0^AT1, tightly packed by residues from extracellular loop 4, TM3 and TM10. Using docking and mutational studies the experiments suggest that MhsT, like SERT, has a second substrate binding site (S2). The authors propose that binding of substrate amino acid tryptophan to this site can occur when the side-chain of Trp33 rotates out of the way. They further propose that allosteric binding of tryptophan could act as a “symport-effector”, facilitating the conformational change of the transporter. This S2 binding site is similar to the JX98 and JX225 binding pocket in B^0^AT1. However, in B^0^AT1 the S2 binding site is clearly inhibitory. It appears likely that binding of JX98 and JX225 to this site prevents closing of the vestibule and the tight packing of amino acid side-chains, which are required for the transition to the inward-facing conformation. We were unable to generate a structure with inhibitor and substrate bound at the same time. This also explains the largely competitive-type of inhibition exerted by JX98 ^10^. In B^0^AT1 the side-chain of Trp56 could act as a pseudo-substrate symport-effector, which would explain why mutation to glycine resulted in a completely inactive transporter, and mutation to leucine resulted in only a small residual transport activity. Notably, tryptophan is a poor substrate of B^0^AT1 with a V_max_ that is 10% of that of the preferred substrate methionine ^14^, which binds to the S1 site ^38^. Thus, tryptophan could displace the side-chain of Trp56, thereby slowing down the transporter by binding as an allosteric inhibitor to the S2 site. Substrates that prefer the S1 site are fast substrates because they do not compete with the side-chain of Trp56.

Overall, our strategy provides a rational path for the design, development, and *in vitro* characterisation of allosteric inhibitors targeting amino acid transporters in the SLC6 family with potential applications for the treatment of various pathologies. Moreover, we provide insight into the role of the allosteric site in the transport mechanism of SLC6 amino acid transporters.

## Acknowledgements

The authors thank Jędrzej Kukułowicz for help with modelling inhibitor binding to B^0^AT1.

## Author contributions

JX, ZH, LD, AY, YJ, AB, and MG performed experiments, analysed data and prepared data for publication, MM, RY, and SB conceived the study, wrote the manuscript and compiled data for publication. All authors edited the manuscript.

## Competing interests

The authors declare no competing interests.

## Data availability

The structures of the ACE2-B^0^AT1 complex bound with JX-98 (PDB: 8WBY, whole map: EMD-37427), and the ACE2-B^0^AT1 complex bound with JX-225 (PDB: 8WBZ, whole map: EMD-37428) have been deposited to the Protein Data Bank (http://wwwrcsb.org) and the Electron Microscopy Data Bank (https://www.ebi.ac.uk/pdbe/emdb/), respectively. The X-ray crystal structure of JX98 and JX225 were deposited in the Cambridge Crystallographic Data Centre (CCDC-2300201 and CCDC-2300200). All other data can be found Mendeley Data, V1, doi: 10.17632/8k89yrd4pz.1

## Notes

### Competing Interest Statement

The authors have declared no competing interest.

## References

1 Broer, S. & Gauthier-Coles, G. Amino Acid Homeostasis in Mammalian Cells with a Focus on Amino Acid Transport. J Nutr 152, 16–28 (2022). 10.1093/jn/nxab342

2 Blau, N., Duran, M., Gibson, K. M. & Dionisi-Vici, C. Physician’s guide to the diagnosis, treatment, and follow-up of inherited metabolic disease. 3–141 (Springer-Verlag, 2014).

3 Holecek, M. Why Are Branched-Chain Amino Acids Increased in Starvation and Diabetes? Nutrients 12 (2020). 10.3390/nu12103087

4 White, P. J. et al. Insulin action, type 2 diabetes, and branched-chain amino acids: A two-way street. Mol Metab, 101261 (2021). 10.1016/j.molmet.2021.101261

5 Palacin, M. & Broer, S. in Physician’s Guide to the Diagnosis, Treatment, and Follow-Up of Inherited Metabolic Diseases (eds B. Thöny, M. Duran, K.M. Gibson, & C. Dionisi-Vici) 85–99 (Springer-Verlag, 2014).

6 Seow, H. F. et al. Hartnup disorder is caused by mutations in the gene encoding the neutral amino acid transporter SLC6A19. Nat Genet 36, 1003–1007 (2004). 10.1038/ng1406

7 Belanger, A. M. et al. Inhibiting neutral amino acid transport for the treatment of phenylketonuria. JCI Insight 3 (2018). 10.1172/jci.insight.121762

8 Belanger, A. J. et al. Excretion of excess nitrogen and increased survival by loss of SLC6A19 in a mouse model of ornithine transcarbamylase deficiency. Journal of Inherited Metabolic Disease n/a (2022). 10.1002/jimd.12568

9 Jiang, Y. et al. Mice lacking neutral amino acid transporter B(0)AT1 (Slc6a19) have elevated levels of FGF21 and GLP-1 and improved glycaemic control. Mol Metab 4, 406–417 (2015). 10.1016/j.molmet.2015.02.003

10 Yadav, A. et al. Novel Chemical Scaffolds to Inhibit the Neutral Amino Acid Transporter B(0)AT1 (SLC6A19), a Potential Target to Treat Metabolic Diseases. Front Pharmacol 11, 140 (2020). 10.3389/fphar.2020.00140

11 Navarro-Garrido, A. et al. Aristolochic acid-induced nephropathy is attenuated in mice lacking the neutral amino acid transporter B(0)AT1 (Slc6a19). Am J Physiol Renal Physiol (2022). 10.1152/ajprenal.00181.2022

12 Pochini, L. et al. Nimesulide binding site in the B0AT1 (SLC6A19) amino acid transporter. Mechanism of inhibition revealed by proteoliposome transport assay and molecular modelling. Biochem Pharmacol 89, 422–430 (2014). 10.1016/j.bcp.2014.03.014

13 Desai, J. et al. Discovery of novel, potent and orally efficacious inhibitor of neutral amino acid transporter B(0)AT1 (SLC6A19). Bioorg Med Chem Lett 53, 128421 (2021). 10.1016/j.bmcl.2021.128421

14 Cheng, Q. et al. Identification of novel inhibitors of the amino acid transporter B(0) AT1 (SLC6A19), a potential target to induce protein restriction and to treat type 2 diabetes. Br J Pharmacol 174, 468–482 (2017). 10.1111/bph.13711

15 Danthi, S. J. et al. Identification and Characterization of Inhibitors of a Neutral Amino Acid Transporter, SLC6A19, Using Two Functional Cell-Based Assays. SLAS Discov, 2472555218794627 (2018). 10.1177/2472555218794627

16 Gerbeth-Kreul, C. et al. A Solid Supported Membrane-Based Technology for Electrophysical Screening of B(0)AT1-Modulating Compounds. SLAS Discov 26, 783–797 (2021). 10.1177/24725552211011180

17 Broer, S. & Gether, U. The solute carrier 6 family of transporters. Br J Pharmacol 167, 256– 278 (2012). 10.1111/j.1476-5381.2012.01975.x

18 Danilczyk, U. et al. Essential role for collectrin in renal amino acid transport. Nature 444, 1088–1091 (2006). 10.1038/nature05475

19 Kowalczuk, S. et al. A protein complex in the brush-border membrane explains a Hartnup disorder allele. FASEB J 22, 2880–2887 (2008). 10.1096/fj.08-107300

20 Yan, R. et al. Structural basis for the recognition of SARS-CoV-2 by full-length human ACE2. Science 367, 1444–1448 (2020). 10.1126/science.abb2762

21 Shi, L., Quick, M., Zhao, Y., Weinstein, H. & Javitch, J. A. The mechanism of a neurotransmitter:sodium symporter--inward release of Na+ and substrate is triggered by substrate in a second binding site. Mol Cell 30, 667–677 (2008). S1097-2765(08)00359-6, 10.1016/j.molcel.2008.05.008

22 Navratna, V. & Gouaux, E. Insights into the mechanism and pharmacology of neurotransmitter sodium symporters. Curr Opin Struct Biol 54, 161–170 (2019). 10.1016/j.sbi.2019.03.011

23 Cheng, M. H. & Bahar, I. Monoamine transporters: structure, intrinsic dynamics and allosteric regulation. Nat Struct Mol Biol 26, 545–556 (2019). 10.1038/s41594-019-0253-7

24 Niello, M., Gradisch, R., Loland, C. J., Stockner, T. & Sitte, H.--H. Allosteric Modulation of Neurotransmitter Transporters as a Therapeutic Strategy. Trends Pharmacol Sci 41, 446–463 (2020). 10.1016/j.tips.2020.04.006

25 Shotwell, M. A., Jayme, D. W., Kilberg, M. S. & Oxender, D. L. Neutral amino acid transport systems in Chinese hamster ovary cells. J Biol Chem 256, 5422–5427 (1981).

26 Broer, S. & Broer, A. Amino acid homeostasis and signalling in mammalian cells and organisms. Biochem J 474, 1935–1963 (2017). 10.1042/BCJ20160822

27 Fairweather, S. J. et al. A GC-MS/Single-Cell Method to Evaluate Membrane Transporter Substrate Specificity and Signaling. Front Mol Biosci 8, 646574 (2021). 10.3389/fmolb.2021.646574

28 Gauthier-Coles, G. et al. Quantitative modelling of amino acid transport and homeostasis in mammalian cells. Nat Commun 12, 5282 (2021). 10.1038/s41467-021-25563-x

29 Wempe, M. F. et al. Metabolism and pharmacokinetic studies of JPH203, an L-amino acid transporter 1 (LAT1) selective compound. Drug Metab Pharmacokinet 27, 155–161 (2012). 10.2133/dmpk.dmpk-11-rg-091

30 Esslinger, C. S., Cybulski, K. A. & Rhoderick, J. F. Ngamma-aryl glutamine analogues as probes of the ASCT2 neutral amino acid transporter binding site. Bioorg Med Chem 13, 1111–1118 (2005). 10.1016/j.bmc.2004.11.028

31 Kuhn, B., Mohr, P. & Stahl, M. Intramolecular hydrogen bonding in medicinal chemistry. J Med Chem 53, 2601–2611 (2010). 10.1021/jm100087s

32 Cavallo, G. et al. The Halogen Bond. Chem Rev 116, 2478–2601 (2016). 10.1021/acs.chemrev.5b00484

33 Cincic, D., Friščić, T. & Jones, W. Isostructural Materials Achieved by Using Structurally Equivalent Donors and Acceptors in Halogen-Bonded Cocrystals. Chemistry – A European Journal 14, 747–753 (2008). 10.1002/chem.200701184

34 Wobst, H. et al. A SMALL MOLECULE SLC6A19 INHIBITOR INCREASES URINARY PHENYLALANINE EXCRETION AND REDUCES ITS PATHOGENIC PLASMA ACCUMULATION IN A PHENYLKETONURIA MOUSE MODEL. Molecular genetics and metabolism. 138, 107502 (2023). 10.1016/j.ymgme.2023.107502

35 Coleman, J. A. & Gouaux, E. Structural basis for recognition of diverse antidepressants by the human serotonin transporter. Nat Struct Mol Biol 25, 170–175 (2018). 10.1038/s41594-018-0026-8

36 Plenge, P. et al. The mechanism of a high-affinity allosteric inhibitor of the serotonin transporter. Nat Commun 11, 1491 (2020). 10.1038/s41467-020-15292-y

37 Quick, M. et al. The LeuT-fold neurotransmitter:sodium symporter MhsT has two substrate sites. Proc Natl Acad Sci U S A 115, E7924–E7931 (2018). 10.1073/pnas.1717444115

38 Li, Y. et al. Structural insight into the substrate recognition and transport mechanism of amino acid transporter complex ACE2-B(0)AT1 and ACE2-SIT1. Cell Discov 9, 93 (2023). 10.1038/s41421-023-00596-2

