## Supplemental data for "Molecular basis of inhibition of the amino acid transporter B^0^AT1 (SLC6A19)"

Supplementary data:

Table S1

Figure S1

Figure S2

Figure S3

Figure S4

Figure S5

Table S2

Table S1: Pharmacological characterisation of compounds by FLIPR and flux assays using leucine as a substrate. Leucine concentration was 150 µM in the flux assay and 1.5 mM in the FLIPR assay. A single dose of 30 µM inhibitor was used for initial screening and measured relative to cinromide (set 0%). A dose-response curve was determined for more potent compounds (n=6).

| Compound | % Activity (FLIPR@30 µM) | IC50 (µM) |
|---|---|---|
| JX8 | 83 ± 4 | n.d. |
| JX42 | 82 ± 7 | n.d. |
| JX43 | 34 ± 4 | 37 ± 33 (Flux) |
| JX55 | 21 ± 2 | 11 ± 3 (Flux) |
| JX63 | 9 ± 2 | 6 ± 0.6 (Flux) |
| JX98 | 38 ± 14 | 7.7 ± 1.9 |

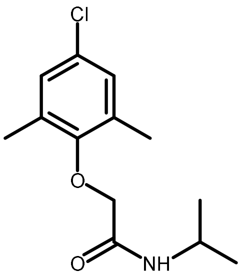

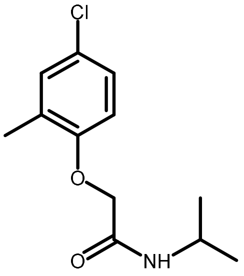

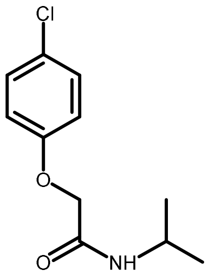

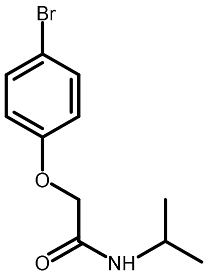

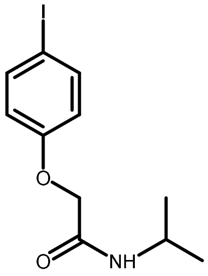

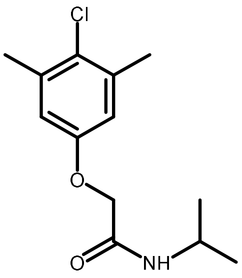

| Compound | % Activity (@30 µM) | IC50 (µM) |
|---|---|---|
| JX108 | 44 ± 6 | 20 ± 10 (Flux) |
| JX109 | 3 ± 1 | 3 ± 0.4 (Flux)<br>0.62 ± 0.07 (FLIPR) |
| JX110 | 11 ± 2 | 4 ± 1.5 (Flux)<br>3.4 ± 1.1 (FLIPR) |
| JX111 | 105 ± 5 | n.d. |
| JX113 | 110 ± 8 | n.d. |
| JX115 | 105 ± 11 | n.d. |
| JX116 | 109 ± 6 | n.d. |

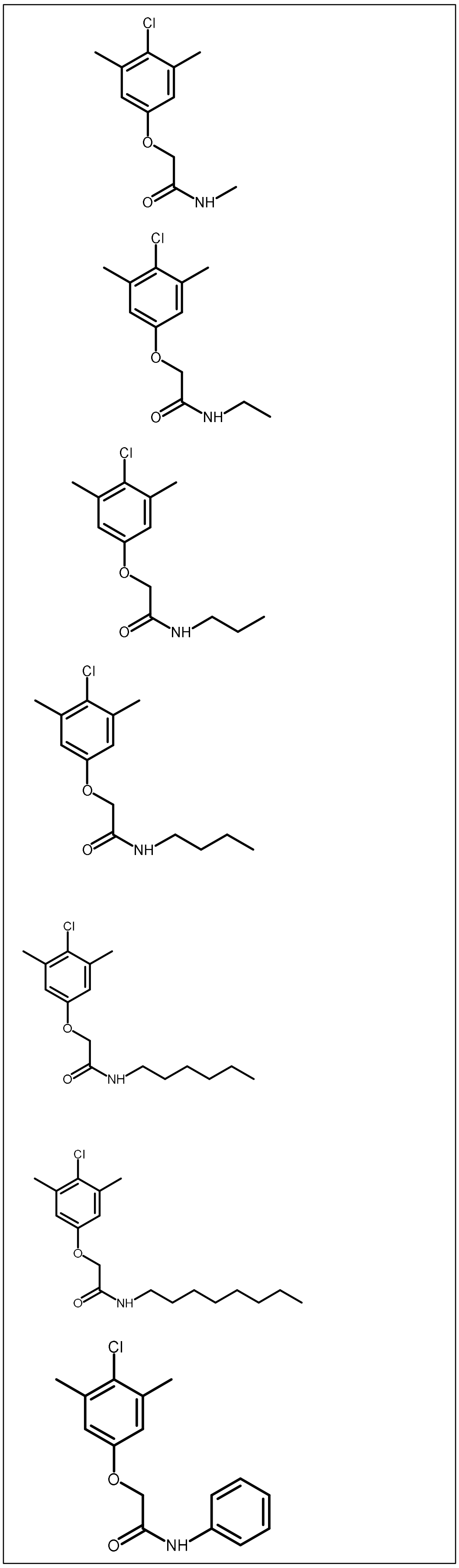

| Compound | % Activity (@30 µM) | IC50 (µM) |
|---|---|---|
| JX117 | 102 ± 10 | n.d. |
| JX118 | 95 ± 6 | n.d. |
| JX127 | 102 ± 10 | n.d. |
| JX128 | 131 ± 10 | n.d. |
| JX129 | 109 ± 10 | n.d. |
| JX130 | 106 ± 13 | n.d. |
| JX131 | 116 ± 4 | n.d. |

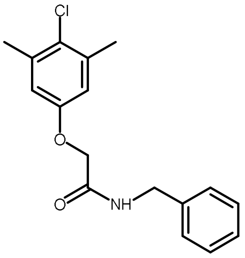

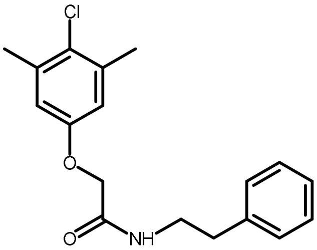

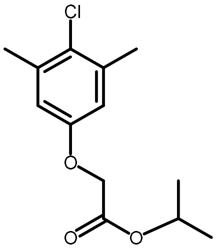

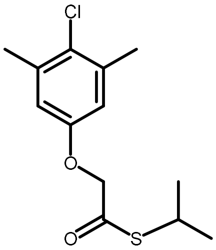

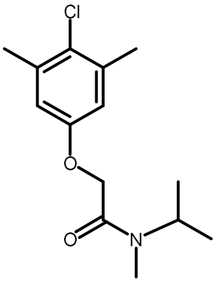

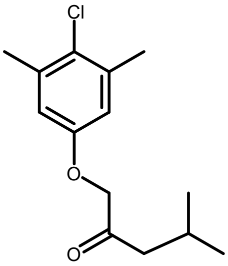

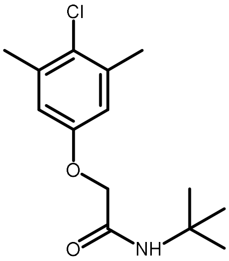

| Compound | % Activity (@30 µM) | IC50 (µM) |
|---|---|---|
| JX132 | 84 ± 11 | n.d. |
| JX133 | 108 ± 12 | n.d. |
| JX147 | 90 ± 10 | n.d. |
| JX148 | 109 ± 10 | n.d. |
| JX149 | 103 ± 10 | n.d. |
| JX160 | 78 ± 9 | n.d. |
| JX161 | 127 ± 9 | n.d. |

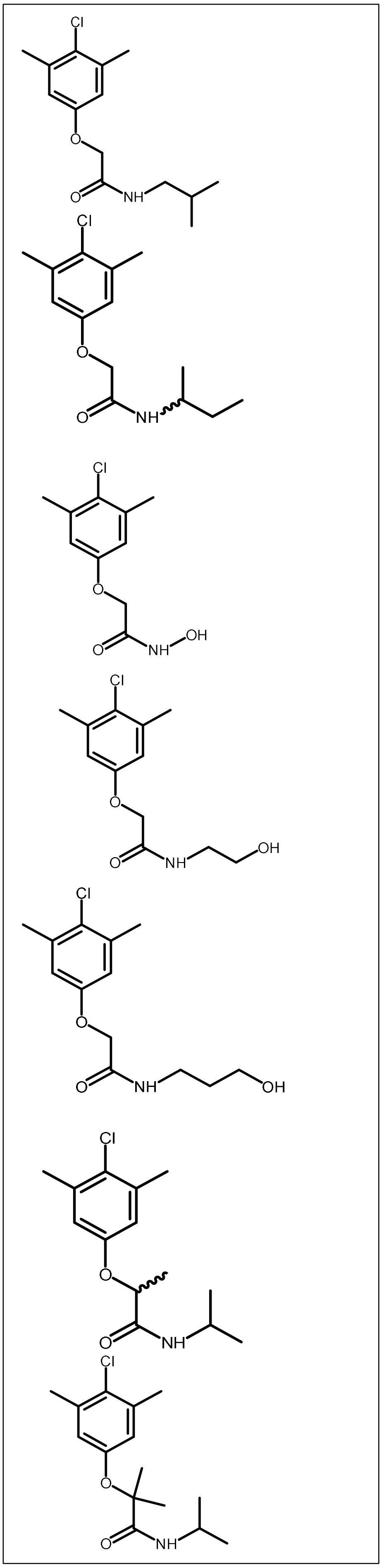

| Compound | % Activity (@30 µM) | IC50 (µM) |
|---|---|---|
| JX171 | 11 ± 2 | n.d. |
| JX172 | 5 ± 1 | n.d. |
| JX173 | 8 ± 5 | 2 ± 0.3 (Flux) |
| JX174 | -3 ± 1 | 1 ± 0.4 (Flux)<br>0.44 ± 0.06 (FLIPR) |
| JX175 | -3 ± 1 | 0.7 ± 0.1 (Flux)<br>0.23 ± 0.06 (FLIPR) |
| JX176 | 10 ± 2 | 6 ± 2 (Flux) |
| JX177 | 1 ± 0.3 | 2 ± 0.4 (Flux) |

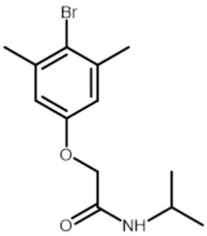

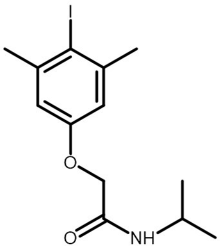

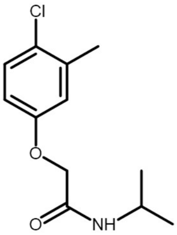

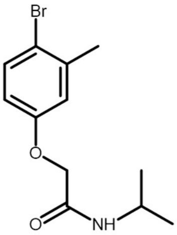

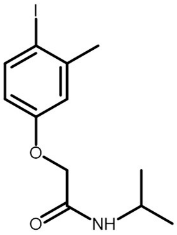

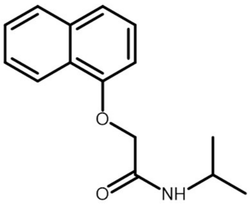

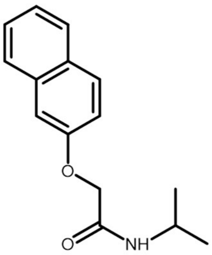

| Compound | % Activity (@30 μM) | IC50 (μM) |
|---|---|---|
| JX184 | -2 ± 1 | 1.2 ± 0.1 (Flux)<br>0.3 ± 0.03 (FLIPR) |
| JX185 | -2 ± 2 | 0.9 ± 0.2 (Flux)<br>0.36 ± 0.02 (FLIPR) |
| JX186 | -3 ± 2 | 0.4 ± 0.1 (Flux)<br>0.099 ± 0.005 (FLIPR) |
| JX189 | 82 ± 11 | n.d. |
| JX191 | 10 ± 4 | n.d. |
| JX192 | 80 ± 22 | n.d. |
| JX193 | 39 ± 11 | n.d. |

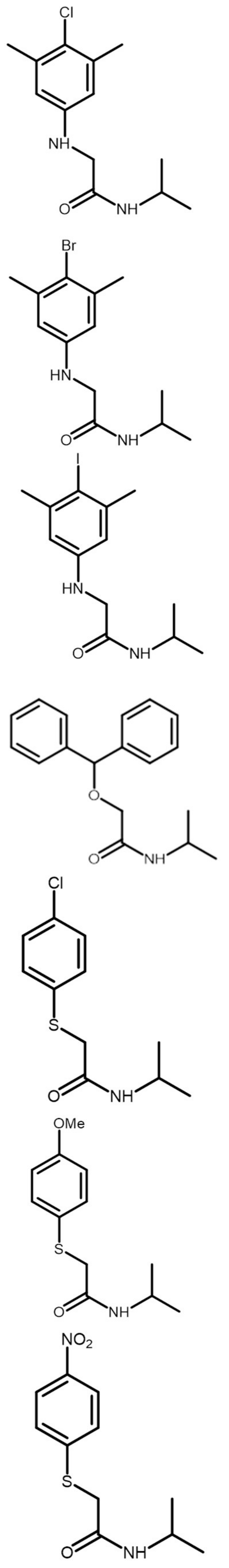

| Compound | % Activity (@30 µM) | IC50 (µM) |
|---|---|---|
| JX225 | n.d. | 0.07 ± 0.01 (FLIPR) |
| JX226 | n.d. | 0.043 ± 0.008 (FLIPR) |
| JX227 | n.d. | 0.081 ± 0.006 (FLIPR) |
| JX228 | n.d. | 0.046 ± 0.006 (FLIPR) |
| JX235 | n.d. | 0.09 ± 0.01 (FLIPR) |
| JX236 | n.d. | 0.04 ± 0.003 (FLIPR) |
| JX237 | n.d. | 0.031 ± 0.002 (FLIPR) |

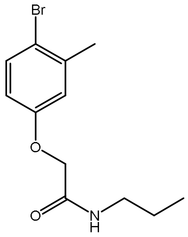

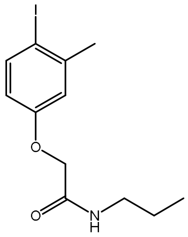

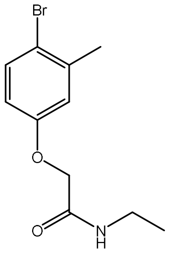

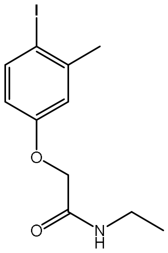

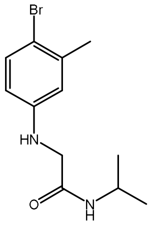

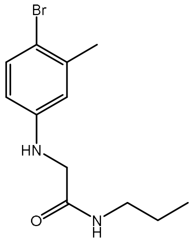

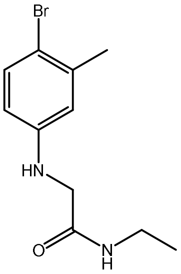

| Compound | % Activity (@30 µM) | IC50 (µM) |
|---|---|---|
| JX238 | n.d. | 0.027 ± 0.005 (FLIPR) |
| JX239 | n.d. | 0.034 ± 0.005 (FLIPR) |
| JX240 | n.d. | 0.033 ± 0.005 (FLIPR) |
| Cinromide | 0 (Reference) | 0.84 ± 1 |

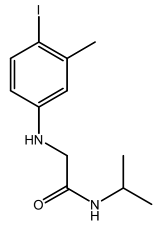

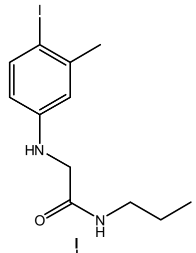

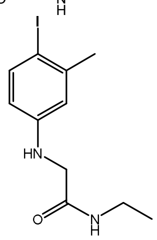

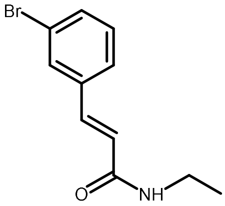

Fig. S1

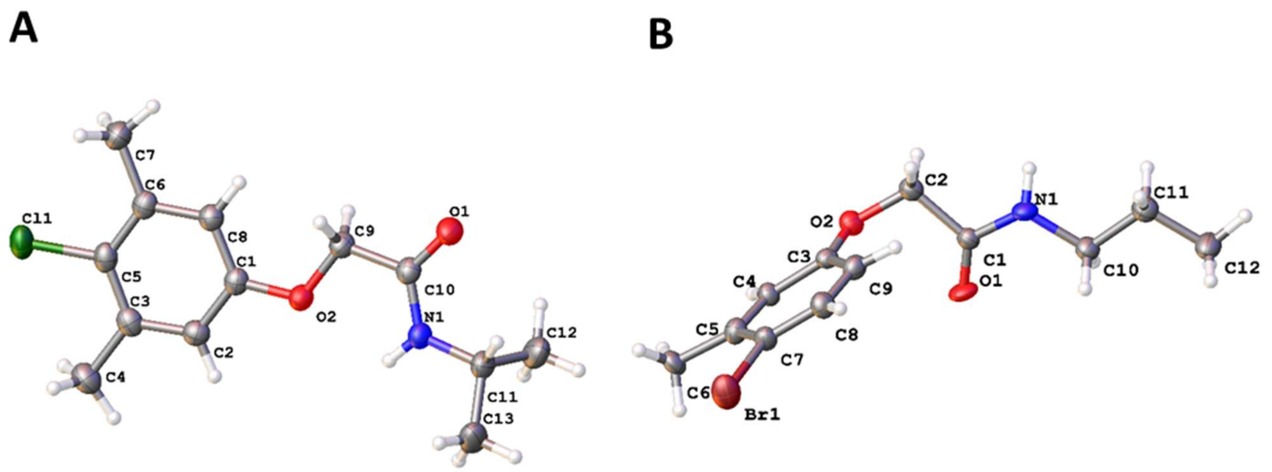

**Figure S1**. Solid state molecular structures of A) JX98, and B) JX225 showing the asymmetric unit (atomic displacement parameters shown at 50% probability level, intra- and intermolecular hydrogen-bonding is omitted).

Fig. S2

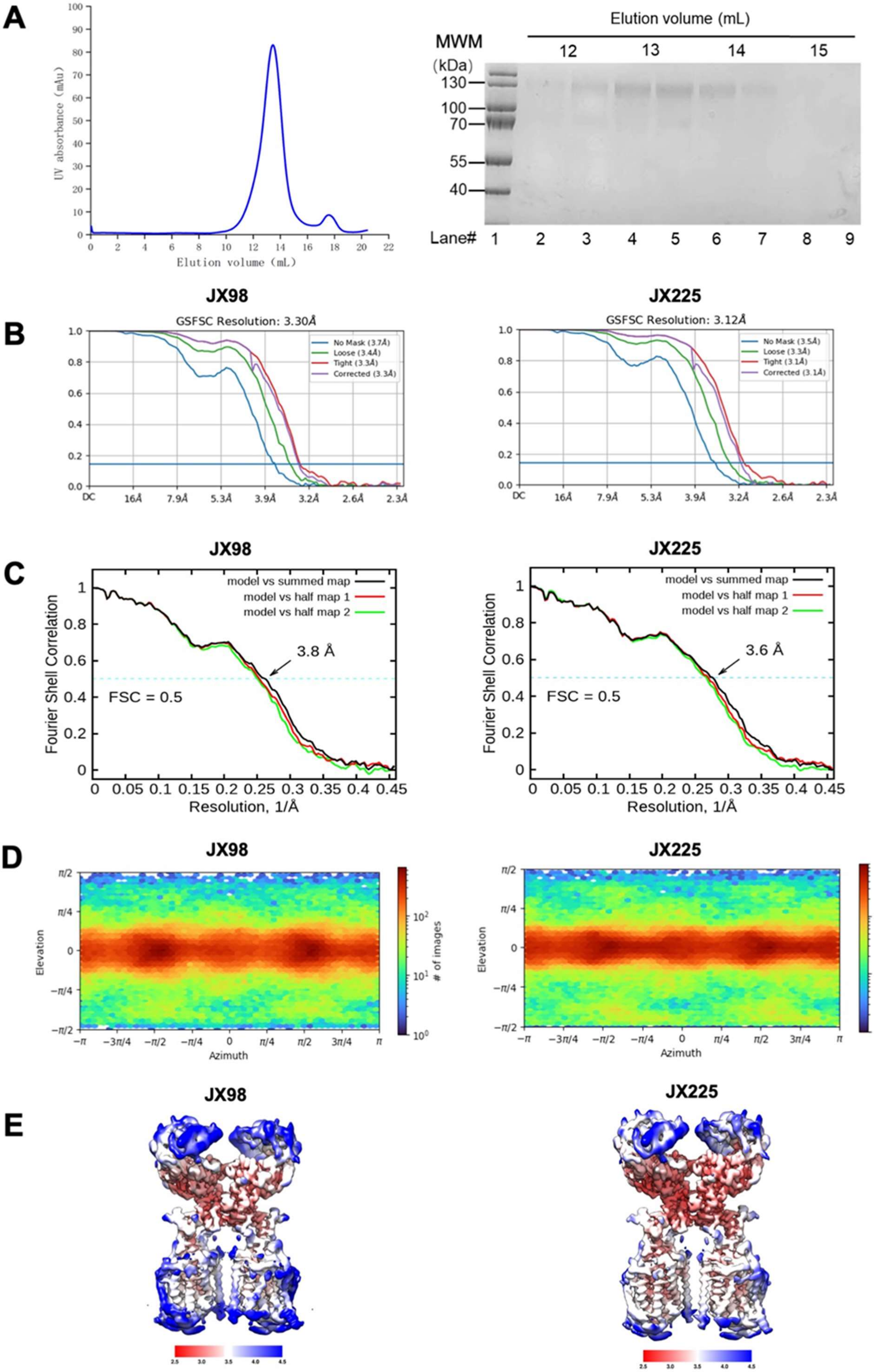
A
UV absorbance (mAu)
Elution volume (mL)
Elution volume (mL)
MWM
(kDa)
12
13
14
15
130
100
70
55
40
Lane# 1 2 3 4 5 6 7 8 9
B
JX98
GSFSC Resolution: 3.30Å
No Mask (3.7Å)
Loose (3.4Å)
Tight (3.3Å)
Corrected (3.3Å)
JX225
GSFSC Resolution: 3.12Å
No Mask (3.5Å)
Loose (3.3Å)
Tight (3.1Å)
Corrected (3.1Å)
DC 16Å 7.9Å 5.3Å 3.9Å 3.2Å 2.6Å 2.3Å
C
JX98
model vs summed map
model vs half map 1
model vs half map 2
3.8 Å
FSC = 0.5
Fourier Shell Correlation
Resolution, 1/Å
JX225
model vs summed map
model vs half map 1
model vs half map 2
3.6 Å
FSC = 0.5
Fourier Shell Correlation
Resolution, 1/Å
D
JX98
JX225
Elevation
Azimuth
### of images
E
JX98
JX225
2.5 3.0 3.5 4.0 4.5

**Figure S2. Cryo-EM reconstruction of ACE2-B$^0$AT1 bound with inhibitors.**

**(A)** Representative size exclusion chromatography (SEC) purification of the ACE2-B$^0$AT1 complex. The protein complex was purified in the presence of GDN. Inset: on the left, the SEC purification diagram of the full-length human ACE2-B$^0$AT1 complex; on the right, SDS-PAGE visualized by Coomassie blue staining. MWM, molecular mass marker. **(B)** Gold-standard Fourier shell correlation (FSC) curves of the ACE2-B$^0$AT1 complex with inhibitors. The resolution was estimated with non-uniform refinement by CryoSPARC 3.1. **(C)** FSC curve of the refined model versus the overall structure of class 2 that it is refined against (black); of the model refined against the first half map versus the same map (red); and of the model refined against the first half map versus the second half map (green). The small difference between the red and green curves indicates that the refinement of the atomic coordinates did not suffer from overfitting. **(D)** Euler angle distribution in the final 3D reconstruction of overall map. **(E)** Cryo-EM density maps colored by local resolution for ACE2-B$^0$AT1 bound with JX98 and JX-225, respectively**.**

Fig. S3

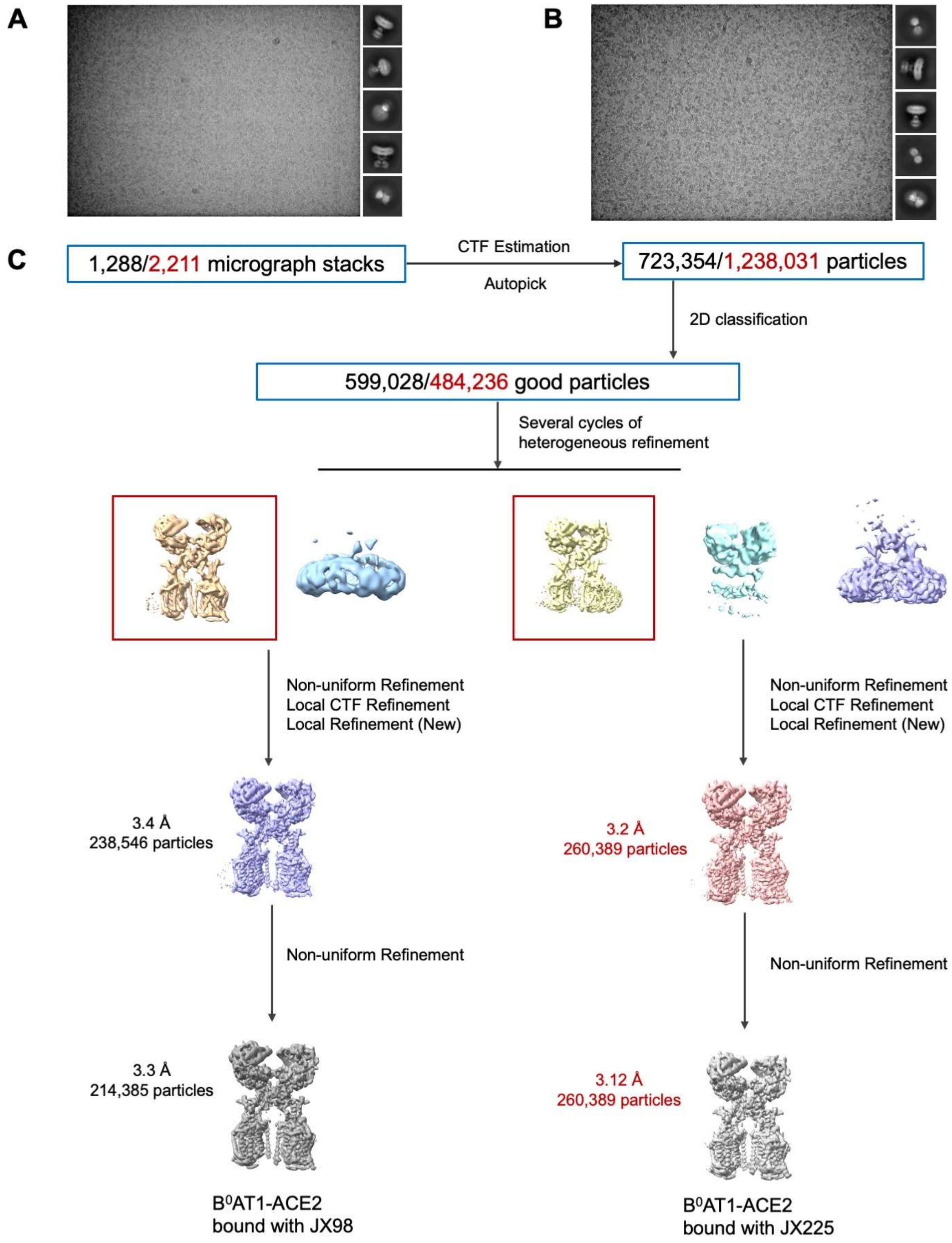

**Figure S3. Flowchart for Cryo-EM data processing.**

Please refer to the 'Data Processing' in Methods section for details.

**Fig. S4**

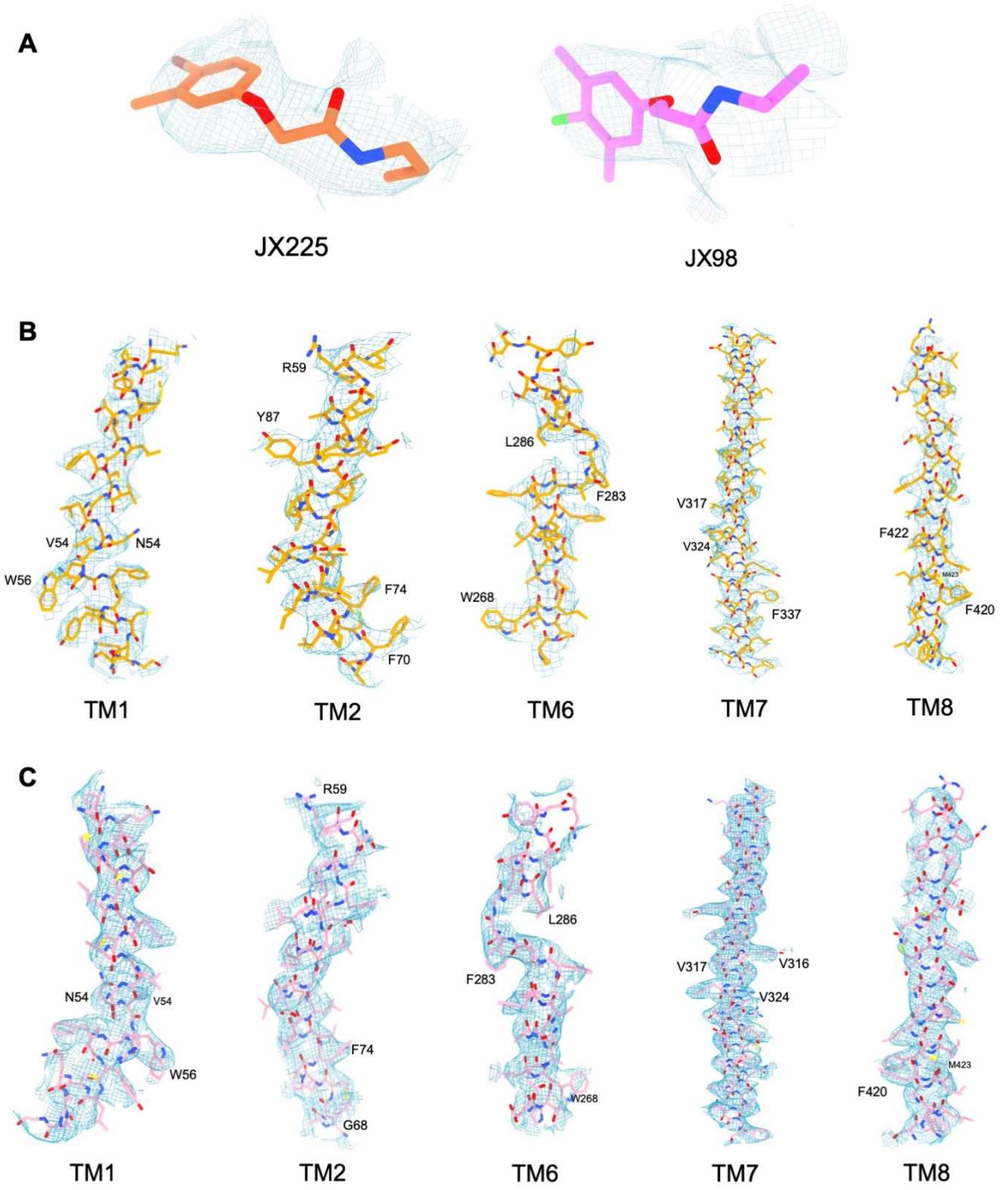

**Figure S4. Cryo-EM density maps of the ACE2-B$^0$AT1 bound with inhibitors.**

**(a-b)** Cryo-EM density maps for the inhibitors bound with B$^0$AT1 shown at counter level of 0.12. (**c-d)** Cryo-EM density maps for the transmembrane helix of B$^0$AT1 shown at counter level of 0.25.

**Fig. S5**

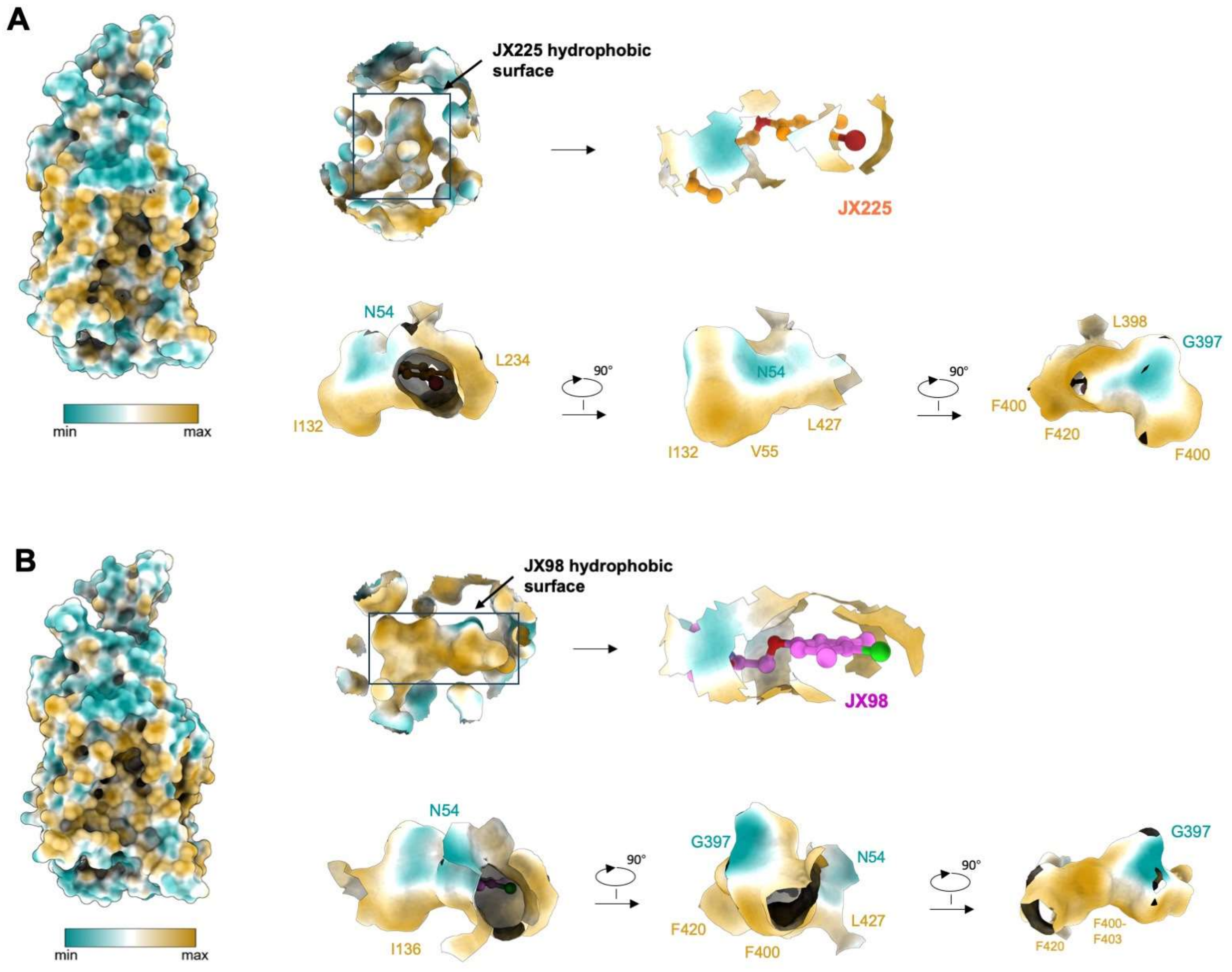

**Figure S5. Hydrophobic surface of the ACE2-B$^0$AT1 bound with inhibitors.**

**(a)** Hydrophobic surface for the JX-225 (coral) bound with B$^0$AT1. (**b)** Hydrophobic surface for the E4 (magenta) bound with B$^0$AT1. Cyan regions represent weak hydrophobicity, while goldenrod areas represent strong hydrophobicity

**Table S2 Cryo-EM data collection, refinement and validation statistics**

| Data collection | | |
|---|---|---|
| EM equipment | Titan Krios (Thermo Fisher Scientific) | |
| Voltage (kV) | 300 | |
| Detector | Gatan K3 Summit | |
| Energy filter | Gatan GIF Quantum, 20 eV slit | |
| Pixel size (Å) | 1.095 | |
| Electron dose (e-/Å2) | 50 | |
| Defocus range (μm) | -1.4 ~ -1.8 | |
| Sample | ACE2-B$^0$AT1 (JX98) | ACE2-B$^0$AT1 (JX225) |
| Number of collected micrographs | 1,288 | 2,211 |
| 3D Reconstruction | | |
| Software | CryoSPARC | |
| Number of used particles (Overall) | 723,354 | 1,238,031 |
| Resolution (Å) | 3.30 | 3.12 |
| FSC threshold for resolution [illegible] | 0.143 | 0.143 |
| Symmetry | C1 | |
| Map sharpening B-factor (Å$^2$) | -90 | |
| Refinement | | |
| Software | Phenix | |
| Model composition | | |
| Protein residues | 2,708 | 2,708 |
| Side chains assigned | 2,708 | 2,708 |
| CC-volume | 0.85 | 0.86 |
| CC-mask | 0.86 | 0.86 |
| B factors (Å$^2$ ) | 143 | 111.6 |
| R.m.s deviations | | |
| Bonds length (Å) | 0.007 | 0.005 |
| Bonds Angle (˚) | 0.732 | 1.073 |
| MolProbitiy score | 1.95 | 2.00 |
| Ramachandran plot statistics (%) | | |
| Preferred | 93.22 | 93.37 |
| Allowed | 6.63 | 6.48 |
| Outlier | 0.15 | 0.15 |
